# BLOC1S1 control of vacuolar organelle fidelity modulates T_H_2 cell immunity and allergy susceptibility

**DOI:** 10.1101/2024.03.21.586144

**Authors:** Rahul Sharma, Kaiyuan Wu, Kim Han, Anna Chiara Russo, Pradeep K. Dagur, Christian A. Combs, Michael N. Sack

**Affiliations:** Laboratory of Mitochondrial Biology and Metabolism, NHLBI, NIH, Maryland, USA; Cardiovascular Branch, NHLBI, NIH, Maryland, USA; Flow Cytometry Core Facility, NHLBI, NIH, Maryland, USA; Light microscopy Core, NHLBI, NIH, Maryland, USA

## Abstract

The levels of biogenesis of lysosome organelles complex 1 subunit 1 (BLOC1S1) control mitochondrial and endolysosome organelle homeostasis and function. Reduced fidelity of these vacuolar organelles is increasingly being recognized as important in instigating cell-autonomous immune cell activation. We reasoned that exploring the role of BLOC1S1 in CD4^+^ T cells, may further advance our understanding of regulatory events linked to mitochondrial and/or endolysosomal function in adaptive immunity. Transcript levels of the canonical transcription factors driving CD4^+^T cell polarization in response to activation showed that, the T_H_2 regulator GATA3 and phosphorylated STAT6 were preferentially induced in BLOC1S1 depleted primary CD4^+^ T (TKO) cells. In parallel, in response to both T cell receptor activation and in response to T_H_2 polarization the levels of IL-4, IL-5 and IL-13 were markedly induced in the absence of BLOC1S1. At the organelle level, mitochondrial DNA leakage evoked cGAS-STING and NF-kB pathway activation with subsequent T_H_2 polarization. The induction of autophagy with rapamycin reduced cytosolic mtDNA and reverses these T_H_2 signatures. Furthermore, genetic knockdown of STING and STING and NF-κB inhibition ameliorated this immune regulatory cascade in TKO cells. Finally, at a functional level, TKO mice displayed increased susceptible to allergic conditions including atopic dermatitis and allergic asthma. In conclusion, BLOC1S1 depletion mediated disruption of mitochondrial integrity to initiate a predominant T_H_2 responsive phenotype via STING-NF-κB driven signaling of the canonical T_H_2 regulatory program.

## INTRODUCTION

The concept that metabolic remodeling is foundational in controlling immune cell fate, function and polarization is now widely established, and termed immunometabolism (1, 2). Here, metabolic substrates modulate immune cell function, for example by diverting mitochondrial metabolism to preferential biosynthetic functions to support immune cell proliferation (3). At the same accumulation or depletion of specific metabolic substrates function as signaling intermediates to regulate immunity via a multitude of mechanisms including: at the level of immune cell chromatin remodeling (4, 5); by transcriptional (6) or posttranslational regulation (7); via intracellular signal transduction (8); via intracellular organelle effects for example by altering mitochondrial fidelity or autophagy (8); and directly through metabolic remodeling (9). Mitochondria themselves, partially stemming from their prokaryote origins, evoke immune activation following the extrusion of intramitochondrial content into the cytoplasm or extracellular space (3). These mitochondrial components are termed damage associated molecular patterns (DAMPs), which when recognized by pattern recognition receptors (PRRs), initiate inflammatory signaling (10). The mitochondrial organelle itself, also functions as a signaling platform, where cytosolic PRRs, RIG-I like receptors (RLRs), binds to the mitochondrial associated viral signaling (MAVS) adaptor protein on the outer mitochondrial membrane to amplify antiviral signaling (11). Although less well characterized, immune-modulatory effects may arise from other vacuolar organelles that regulate metabolism, including autophagosomes and the endosome-lysosome system (12–14). We reasoned that the study of intracellular vacuolar organelle regulatory control mechanisms may enhance our understanding of the role of this organelle biology in immune cell activation.

To interrogate this further, we proposed to focus on a candidate protein, BLOC1S1/GCN5L1, which is emerging as an important regulator of mitochondrial and of endo- lysosome homeostasis (15). This pleotropic protein modulates mitochondrial turnover (16, 17) and metabolic function (18, 19), and controls key aspects of autophagosome (17), endosome (20) and lysosome trafficking (21, 22), recycling (23, 24), and function (15, 25, 26). These diverse effects stem from the role of BLOC1S1 as an interacting cofactor that modulates mitochondrial and cytosolic protein acetylation and binds to, and regulates, the function of cytoskeletal and molecular motor proteins. Hence, despite the importance of mitochondrial function in immune modulation and the role of endo-lysosomal biology in antigen presentation, cytokine release and in the control of intracellular pathogens, the role of BLOC1S1 in the immune system has to our knowledge not been previously investigated.

To explore this, conditional knockout *bloc1s1* mice were generated using the CD4-Cre- recombinase, employing a similarly approach to the prior deletion of BLOC1S1 in other cell types including the heart (27) and liver (28). Conditional CD4^+^ T cell knockout mice (TKO) were viable and showed robust depletion of BLOC1S1 levels. In this paper, we demonstrate that activated TKO CD4^+^ T_0_ cells show significantly higher IL-4, IL-5, and IL-13 production, a greater propensity to T_H_2 differentiation and that they had elevated phosphorylation of NF-κB, STAT6 and STING. Moreover, TKO mice were highly susceptible to atopic dermatitis (AD) with a marked eosinophilic infiltrate and produced significantly more type 2 cytokines relative to control animals. In addition, OVA-sensitized TKO mice had significantly increased serum IgE and airway inflammation after OVA challenge. Our data suggest a novel role for BLOC1S1 in controlling CD4^+^ lineage commitment and type 2 allergic responses.

## MATERIALS AND METHODS

### Mice

The NHLBI Animal Care and Use Committee approved all animal studies used in this protocol. The mice were maintained on a 12-h light/dark cycle and housed 3-5 mice per cage with free access to water and normal chow diet (LabDiet, 5001). BLOC1S1 CD4^+^ T cell knockout (TKO) mice were generated by crossing BLOC1S1^flox/flox^ mice with CD4-Cre-recombinase mice, as we had previously. All mice were generated in the C56BL/6b background. All experiments used 8-12 week old C57BL/6^flox/flox^ (control) and CD4^+^ TKO mice (backcrossed > 10 generations).

### Mouse CD4^+^ T cell isolation and cytokine assay

All in vitro assays were performed using between three and five mice per group. CD4^+^ T cells were negatively selected from the spleenocytes using CD4^+^ T cell isolation kit (Miltenyi Biotec) and cultured in RPMI 1640 media supplemented with 25 mM HEPES, 10% FBS, and Penicillin/Streptomycin. Mice CD4^+^ T cells (4x10^5^/well in 96-well plate) were activated with plate-coated αCD3 (5 µg/ml, Biolegend) and αCD28 (10 µg/ml, Biolegend) for 3 days. Also, CD4^+^ T cells (4X10^5^/well in 96-well plate) were differentiated into T_H_2 T cell subtype by incubation with specific supplement for T_H_2 differentiation (mouse IL-2, mouse IL-4 and rat anti mouse IFNγ with 1:100 dilution (STEMCELL Technologies)) and incubated for 3 days on plates coated with aCD3 and aCD28 antibodies. Supernatants were collected, centrifuged to remove cells and debris, and stored at -80^0^ C. The levels of cytokines, including IFNγ, TNFα, IL-4, IL-5, IL-13, IL-10, and IL-17 were measured by ELISA (R&D systems). Results were normalized to cell number using CyQuant cell proliferation assay (Invitrogen) or BCA protein assay (Pierce).

### Human CD4^+^ T cell isolation and cytokine assay

Primary peripheral blood mononuclear cells (PBMCs) were isolated from human blood by density centrifugation using Lymphocyte Separation Medium (MP Biomedicals). CD4+ T cells were negatively selected from PBMCs using CD4^+^ T cell isolation kit (Miltenyi Biotec) and cultured in RPMI 1640 media supplemented with 25 mM HEPES, 10% FBS, and Penicillin/Streptomycin. Human CD4^+^ T cells (4X10^5^/well in 96-well plate) were activated with plate-coated αCD3 (5µg/ml, Biolegend) and αCD28 (10µg/ml, Biolegend) for 3 days. Supernatants were collected, centrifuged to remove cells and debris, and stored at -80^0^ C. The levels of cytokines, including IL-4, IL-5 and IL-13 were measured by ELISA (R&D systems). Results were normalized to cell number using CyQuant cell proliferation assay (Invitrogen) or BCA protein assay (Pierce).

### RNA Isolation and Quantitative PCR (qRT-PCR) analysis

Total RNA was extracted using NucleoSpin RNA kit (Macherey-Nagel) and cDNA was synthesized with the SuperScript III First-Strand Synthesis System for RT-PCR (Thermo Fischer Scientific). Quantitative real-time PCR was performed using FastStart Universal SYBR Green master (Roche) and run on LightCycler 96 Systems (Roche). Relative gene expression was quantified by normalizing cycle threshold values with 18S rRNA using the 2^-ΔΔCt^ cycle threshold method. To measure mitochondrial DNA (mtDNA) in the cytosol of CD4^+^ T cells, 8 × 10^6^ cells were homogenized with a Dounce homogenizer in 10 mM Tris solution (pH 7.4), containing 0.25 M sucrose, 25 mM KCl, 5 mM MgCl_2_ and protease inhibitor, and then centrifuged at 700 × g for 10 min at 4°C. Cytosolic fractions were prepared by centrifugation at 10,000 × g for 30 min at 4°C and DNA was isolated from them using the DNeasy Blood & Tissue kit (Qiagen). The copy number of mitochondrial DNA encoding 16S RNA (RNR2) and non-coding D-loop region was measured by quantitative real-time qRT-PCR. Primer sequences are provided in Supplementary Table 1.

### Immunoblot Analysis

Mice or Human CD4 T cells were lysed using RIPA buffer supplemented with protease inhibitor cocktail (Roche) and phosphatase inhibitors (Pierce). Lysates were separated by NuPAGE 4- 12 % Bis-Tris Gels (Thermo Fischer Scientific) and transferred to nitrocellulose membranes (Trans-Blot Turbo Transfer Systems (Bio-Rad Laboratories). Membranes were blocked with Odyssey Blocking Buffer (Li-Cor) and incubated with appropriate antibodies overnight at 4^0^ C. List of Primary antibodies used are provided in Supplementary Table 2. The secondary antibody conjugated with IRDye 800 CW or IRDye 680RD (Li-Cor) were then incubated for 1 hour at room temperature. Immunoblots were scanned using an Odyssey Clx imaging system (Li-Cor Biosciences). Protein band intensity was quantified using ImageJ software (National institute of health).

### Flow cytometry for cell phenotyping

Activated TKO CD4^+^ T cells were activated with Cell stimulation cocktail plus protein transport inhibitors (eBioscience) and PMA (500 ng/ml, Sigma) for 4 hrs and then incubated with antibodies targeting cell surface markers, transcription factors and (BD, Biolegend). Data were acquired with FACSymphony (BD) and post-acquisition analysis was performed using Flowjo 9.9.6 (Treestar Inc.). Analysis excluded debris and doublets using light scatter measurements, and dead cells by live/dead stain. Gating strategies used to identify immune cell subsets are provided in Supplementary Table 3. Briefly, the cells were first gated for singlets (FSC-H *vs*. FSC-A) and further analyzed for their uptake of the Live/Dead Zombi violet stain (Biolegend) to determine live versus dead cells in CD3^+^CD4^+^. The expression of transcription factors and cytokines is then determined for T cell polarization within this gated population.

### Characterization of BLOC1S1 signaling in primary human T cells

Primary CD4^+^ T cells were cultured in RPMI with 1% FBS on αCD3/αCD28 antibody-coated plates for 3 days. The following inhibitors were added to the cells for 24 hours before harvesting: the NF-κB inhibitor JSH23 (2 µM, Tocris Bioscience), STING inhibitor H151 (5 µM, Tocris Bioscience), or Rapamycin (2 µM, Selleckchem). Cytotoxicity was assessed using the CyQUANT LDH Cytotoxicity Assay (Invitrogen). On the third day, cells were centrifuged for Western blot or RNA analysis and the supernatant was collected for ELISA assay.

### Genetic knockdown experiments

For the siRNA knockdown experiments, primary CD4^+^ T cells were transfected with 1.5 µM SMARTpool Accell BLOC1Sa and STING siRNA or Accell control siRNA in Accell siRNA delivery medium (Dharmacon). Knockdown cells were activated on αCD3/αCD28 (Biolegend) antibody-coated plates for 3 days.

### Experimental atopic dermatitis

Calcipotriol (MC903:Sigma Aldrich) was resuspended in ethanol at 50 µM. A total of 1 nM was applied daily to the outer and inner surfaces of the left ear (20 ml/ear) as described previously (29) for 15 days, as follow: 5 days topical treatment, 2 days interruption, 5 days treatment, 2 days interruption, 1 day topical treatment. Ethanol (20 ml/ear) was applied to the contralateral ear as the vehicle control. Auricular lymph node of the MC903 or vehicle-treated mice were extracted at the end of the study and ear sections were either fixed in 10% Formalin for histology or stored at -80^0^ C. CD4^+^ T cells were negatively selected from the minced auricular lymph nodes using the CD4^+^ T cell isolation kit (Miltenyi Biotec) and cultured in RPMI 1640 media supplemented with 25 mM HEPES, 10% FBS, and Penicillin/Streptomycin. Auricular lymph node CD4^+^ T cells (2x10^5^/well in 96-well plate) were activated with plate-coated αCD3 and αCD28 for 3 days. Supernatants were collected, centrifuged to remove cells and debris, and stored at -80 ^0^C.

### Experimental allergic airway inflammation

Mice were sensitized with ovalbumin (OVA, MedChem Express) to induce allergic airway inflammation (30). Mice were administered 20 mg OVA in 4mg alum hydroxide (InvivoGen) by intraperitoneal (i.p) injection on days 0 and 7 and subjected to airway exposure with 40µg OVA in PBS on day 11-14. 24 h after the last OVA aerosol challenge, lungs from allergen sensitized and challenged mice were taken and a section was fixed in 4% paraformaldehyde (PFA) for histology and the residual lung was used to isolate CD4^+^ T cells. List of reagents provided in supplementary table 4.

### Histology

Lungs from the allergen-sensitized and challenged mice experiment and ears from the MC903 and ethanol vehicle treated mice experiment were fixed in 4% PFA for histology. The fixed samples were processed and stained with hematoxylin and eosin (H&E) and Ki67 immunohistochemical staining (Histoserv).

### Immunofluorescence staining and microscopy

CD4+ T cells were activated as earlier described and adhered to glass slides coated with poly-L-lysine (Sigma Aldrich). The cells were fixed with 4% paraformaldehyde for 15 min at room temperature. After the cells were washed three times with PBS, cells were blocked in 5% BSA and 0.1% Triton X for 1 hour at room temperature. Cells were then incubated with appropriate primary antibody overnight at 4^0^C. Then, the cells were washed three times with PBS and were incubated with the appropriate Alexa secondary Abs for 60 minute at room temperature in the dark. Nuclei were counterstained blue with DAPI. All fluorescent imaging performed using a Zeiss 880 confocal microscope and a Plan-Apochromat 63x(1.4 N.A.). Four color images of DAPI, Alexa 488, Alexa 561, and Alexa 633 were collected using 405nm, 488nm, 561nm and 633nm with emission bandwidths of 415-480nm, 490-556nm, 565-659nm and 641-735nm respectively. Three color imaging of DAPI, Alexa 488, and Alexa 561 were collected using 405nm, 488nm, and 561nm excitation with 415-481nm, 480-569nm, and 570- 709nm, respectively. Pixels sizes varied from 0.68-0.98 microns. The pinhole was set to 1A.U. for all experiments and z-stacks were taken with an interslice spacing of 300nm. For experiments where intensity was compared between treatments laser excitation power did not vary more than 0.1%. Images were deconvolved assuming an idealized point spread function using the Hyugens software program (SVI, Hilversum, Netherlands)..

### Quantification of confocal Immunofluorescent images

Immunofluorescent images were quantified using the open-source program FIJI (31). In short, regions of interest (ROI’s) were manually drawn on each cell to segment the nucleus from the cytoplasm. From the segmented images summed intensity from each compartment across z-stacks was then calculated.

### Statistical analysis

Graphs were plotted and analyzed using GraphPad Prism 9. Statistical analysis was performed with either a two-tailed unpaired student t test (for paired data) or two-way ANOVA with Tukey’s post hoc test for experiments with multiple groups. Probability values of <0.05 were considered statistically significant. Data are shown as mean ± SEM. Asterisks denote p value (*p<0.05, **p<0.01, ***p<0.001 and ****p<0.0001).

## RESULTS

### BLOC1S1 depleted CD4^+^ T cell preferentially augments T_H_2 immune cell responsiveness

The approach to generate CD4^+^ T cell-specific BLOC1S1 knockout (TKO) mice is depicted in Supplemental Figure 1A. Separation of CD4^+^ cells from the residual splenic pool (CD4- cells) as depicted in Supplemental Figure 1B, with subsequent qRT-PCR shows robust reduction in bloc1s1 transcript levels in the CD4^+^ pool compared to the Lox-P littermate control mice (Supplemental Figure 1C). To initially characterize CD4^+^ T cell immunoresponsiveness, primary CD4^+^ T cells were activated by antibodies directed against CD3 and CD28 to engage the T cells receptors (TCRs). ELISA assays to assess cytokine secretion showed that TCR engagement in TKO cells significantly increased levels of interleukins (ILs) 4, 5, 10 and 13, reduced IL-17 and without effects on interferon gamma (IFN-γ) and TNFα compared to control cells (Figure 1A). We then assayed the transcript levels of canonical transcription factors driving CD4^+^ T cell polarization in response to TCR activation. The expression of *tbet* (Tbx21 - T_H_1 polarizing transcription factor (TF)), *rorc* (RAR related orphan receptor C – Th17 polarizing TF) and *foxp3* (forkhead TF family member p3 – Treg polarizing TF) were not different between genotypes in T_H_0 cells (Supplemental Figure 1D). In contrast, the gene encoding GATA3, and its cognate protein level were markedly induced in TKO cells (Figure 1B-D). Flow cytometry analysis validated the increase in GATA3^+^,IL4^+^ in TKO CD4^+^ T cells (Figure 1E) with no changes in TBET^+^,IFNg+ or Rorc^+^,IL17^+^ cells (Supplemantal Figure 1E). The gating profiles of CD4^+^ T cells for flow cytometry is shown in Supplemental Figure 1F. We then assessed the effect T_H_2 polarization on the secretion of canonical T_H_2 cytokines. Consistent with the T_H_0 data, levels of IL-4, IL-5 and IL-13 secretion were induced to a greater extent in the TKO cells (Figure 1F). To further validate this response to diminished BLOC1S1 levels siRNA targeting bloc1s1 or scrambled constructs were transfected into primary human CD4^+^ T cells. The efficiency of Bloc1s1 knockdown (KD) was ≈ 80% and transcript levels of GATA3 were induced coordinately (Supplemental Figure 1G). In parallel, transcript levels and secreted T_H_2 cytokines were induced to a greater extent following bloc1s1 knockdown (Supplemental Figures G-H). As GATA3 functions as the master regulator driving T_H_2 differentiation and given the effect of reduced BLOC1S1 on T_H_2 cytokine release, this manuscript subsequently focused on the study of BLOC1S1 effects on T_H_2 biology.

**Figure 1.**
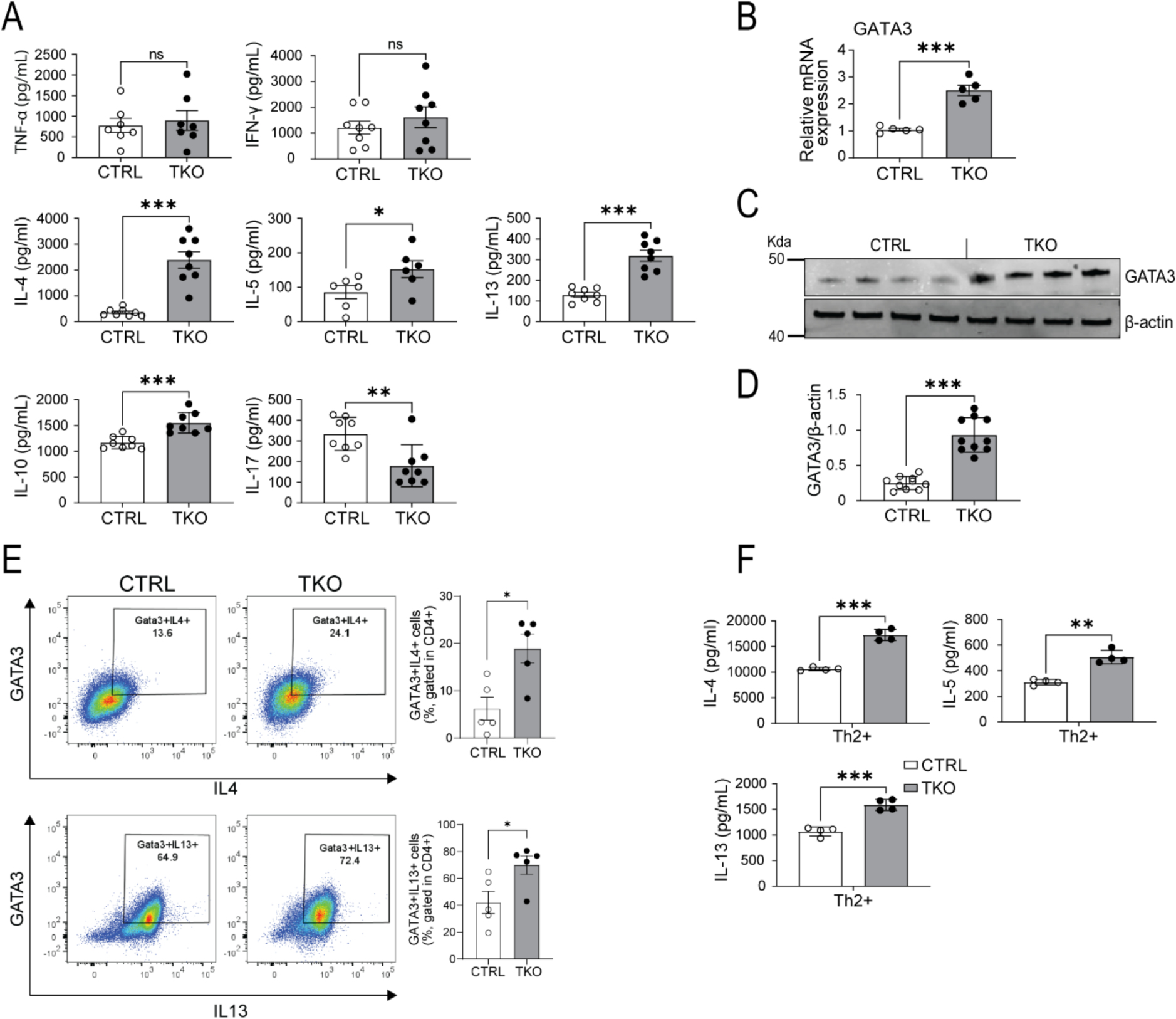
BLOC1S1 depleted CD4^+^ preferentially augments T_H_2 immune responsiveness. **(A)** TH1, T_H_2, Treg and TH17 associated cytokines released from CD4^+^ T cells isolated from the spleen of control (CTRL) and TKO mice, activated with antibodies directed against CD3 and CD28 for 3 days (n=7-8 per group). **(B)** qRT-PCR showing relative mRNA expression levels of GATA3 (n=5 per group). **(C)** Representative immunoblot analysis of GATA3 and β-actin. **(D)** Densitometry analysis of the relative protein levels of GATA3/β-actin in CD4^+^ T cell lysate from spleen of control and TKO mice (n=10 per group). **(E)** Representative flow-cytometric analysis of intracellular cytokines GATA3^+^IL4^+^ and GATA3^+^IL13^+^ in CD4^+^ T cells (n=5 per group). Values represent mean ± SEM.. **(F)** IL-4, IL-5 and IL-13 cytokine release in CD4^+^ isolated from the spleen of control and TKO mice, activated with αCD3 and αCD28, supplemented with T_H_2 differentiation cocktail for 3-4 days (n=4 per group). Values represent mean ± SEM. *P<0.05, *p<0.05, **p<0.01, ***p<0.001 vs. control mice using unpaired two- tailed student-t-test. FSC, forward scatter; SSC, side scatter.

### Enhanced phosphorylation of IKK, NF-κB and STAT6 and GATA3 protein levels in TKO CD4^+^ T cells

The transcriptional and signaling pathways underpinning T_H_2 lineage polarization are well characterized, with GATA3 and STAT6 playing central roles (32). More recently in the context of allergic asthma, NF-κB signaling has been implicated in GATA3 induction and subsequent T_H_2 polarization (30). We therefore explored the activity of these regulatory molecules. To investigate these pathways splenic CD4^+^ T cells were isolated from control and TKO mice and exposed to TCR engagement. In activated T_H_0 cells, in the absence of BLOC1S1, phosphorylation and activation of IκB, NF-κB and STAT6 were markedly induced, in parallel with the induction of GATA3 (Figure 2A-B). In support of the functional contribution of this NF-κB - GATA3 pathway, pharmacological inhibition of NF-κB with JSH23 resulted in diminished NF-κB phosphorylation and GATA3 steady-state levels relative to DMSO treated controls (Figure 2C-D). In parallel, JSH23 blunted T_H_2 cytokine induction to a greater extent in the TKO T_H_0 cells (Figure 2E).

**Figure 2.**
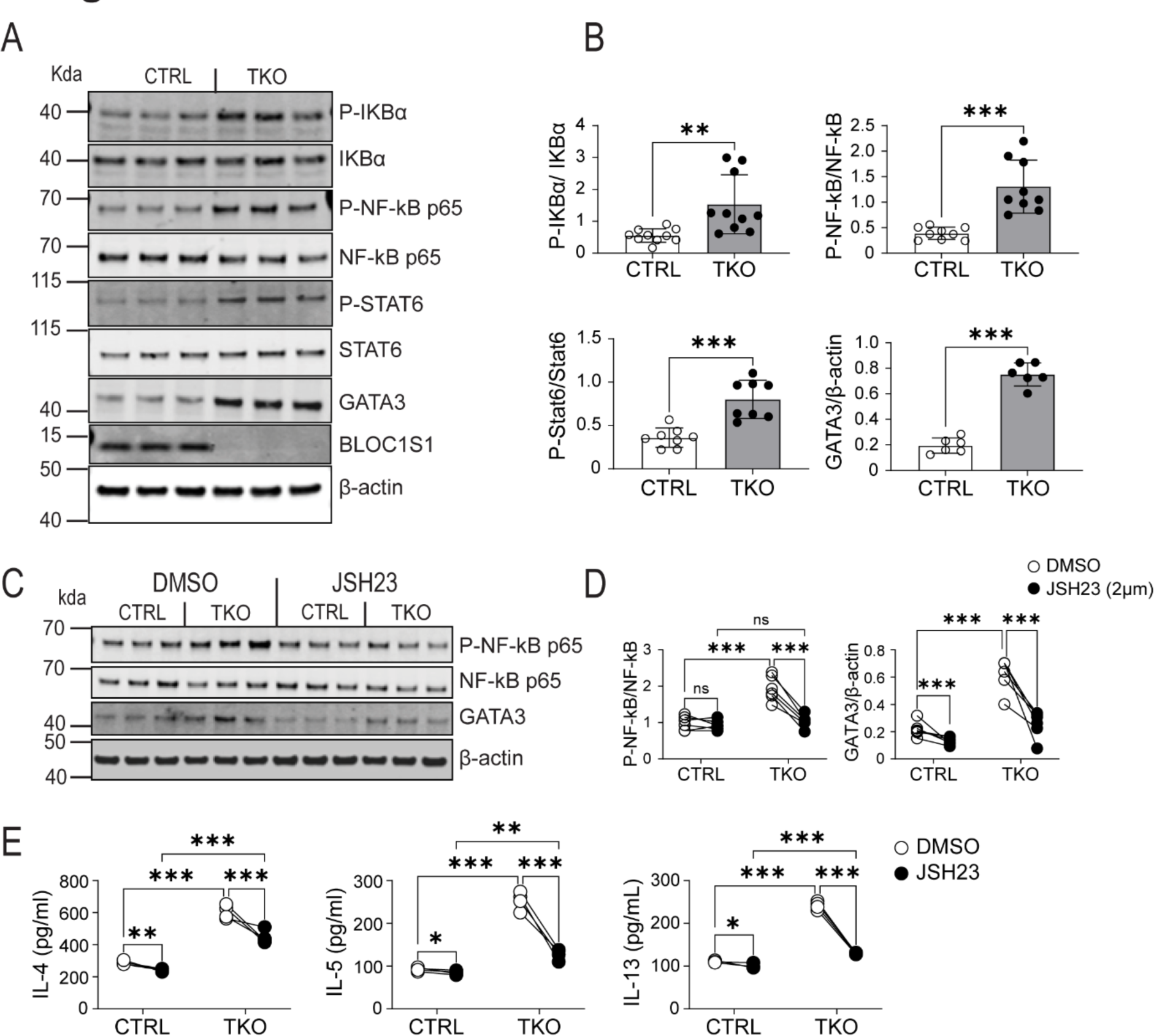
BLOC1S1 deficiency results in enhanced IKK, NF-kB and STAT6 phosphorylation and GATA3 activity. **(A)** Representative immunoblot for phospho-IκBα, IκBα, phospho-NF-κB p65, NF-κB p65, phospho- STAT6, STAT6, GATA3, BLOC1S1 and β-actin from splenic CD4^+^ T cells lysates from control and TKO mice. **(B)** Quantitation of the ratio of P-IκBα/IκBα, P-NF-κB p65/NF-κB p65, P-STAT6/STAT6 and GATA3/β-actin by densitometry analysis (n=6-10). **(C)** Representative immunoblot analysis of P- NF-κB P65, NF-κB p65, GATA3 and β-actin from contron and TKO CD4^+^ T cells incubated with either DMSO or 2 μM JSH23 for 12 hours. **(D)** Quantitation of the ratio of P-NF-κB p65/NF-κB p65 and GATA3/β-actin by densitometry analysis (n=5-6). **(E)** IL-4, IL-5 and IL-13 cytokine release in activated CD4^+^ T cells incubated with DMSO or JSH23 2uM for 12 hours (n=5-6). Values represent mean ± SEM. *p<0.05, **p<0.01, ***p<0.001 vs control mice by two-way ANOVA followed by the Tukey’s post hoc test or unpaired two-tailed student-t-test.

### Vacuolar organelle perturbations drive BLOC1S1 accentuated T_H_2 immunity via STING induction

Interestingly, nucleic acid sensing has been established to drive T_H_2 rather than T_H_1 or T_H_17 differentiation (33) and mitochondrial DNA (mtDNA) release activates the double-stranded nucleic acid sensing cGAS-STING PRR pathway (34). Considering the role of BLOC1S1 in sustaining mitochondrial fidelity (15), we investigated this as a putative mechanism whereby BLOC1S1 depletion regulates T_H_2 signaling. The amount of cytosolic mtDNA, as measured using qRT-PCR showed higher levels of the mitochondrial encoded 16S RNA (RNR2) and non-coding D-loop region of mtDNA in the cytosol in TKO cells (Figure 3A). Given this, it was unsurprisingly that the levels of cGAS and extent of STING phosphorylation, as measured by immunoblot analysis, were induced in TKO CD4^+^ T cells (Figure 3B-C). To validate this, confocal microscopy using fixed CD4^+^ T cells were labelled with fluorescent-tagged antibodies to endogenous STING in parallel with nuclear DNA labeling with DAPI. Here too, STING levels were confirmed to be elevated in the TKO cells (Figure 3D-E). In parallel, fluorescent-tagged antibodies targeting cGAS and cytosolic dsDNA (35), similalry showed their induction and greater cytosolic colocalization in TKO cells (Figure 3F-G). As BLOC1S1 deficiency is known to result in autophagosome accumulation (24) and evident here by increased LAMP1 and LC3-II in TKO cells (Figure 3B-C and Supplemental Figure 2), we assessed if driving autophagy with rapamycin may ameliorate this inflammatory program. Here the inhibition of mTORC1 with rapamycin blunted LAMP1 levels in parallel with a further increase in LC3-II levels in TKO cells, suggesting induction of autophagy (Figure 3H-I). In parallel, rapamycin reduced cytosolic levels of mtDNA in control and TKO cells (Figure 3J) and blunted IL-4, IL-5 and IL-13 secretion to a greater extent in the TKO cells (Figure 3K).

**Figure 3.**
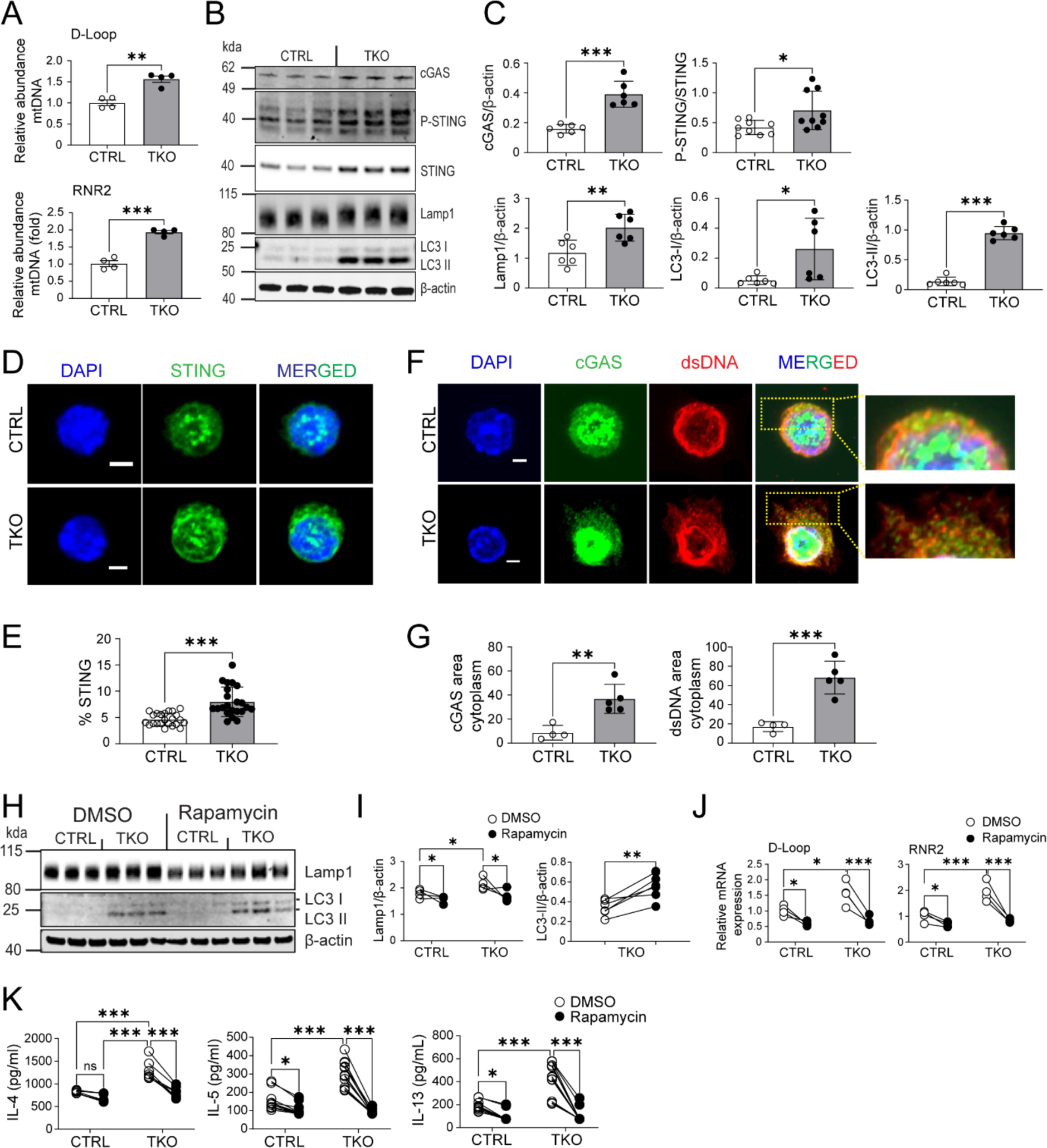
BLOC1S1 deficiency results in mtDNA release into cytosol and activation of cGAS- STING pathway. **(A)** qRT-PCR showing relative mRNA expression levels of D-Loop and RNR2 in CD4^+^ T cells (n=4). **(B)** Representative immunoblot of cGAS, P-STING, STING, Lamp1, LC3 I/II and β-actin from CTRL and TKO CD4^+^ T cells. **(C)** Protein quantitation and ratio of cGAS/β-actin, P-STING/STING, Lamp1/β- actin, LC3 I/β-actin, LC3 II/β-actin and GATA3/β-actin by densitometry analysis (n=6-9). **(D)** Representative fluorescence images of STING (green) in activated CD4^+^ T cells. Nuclei were counterstained with 4’,6-diamidino-2-phenylinodole (DAPI) ((blue). Scale bar = 2 μm. **(E)** Semiquantitative analysis of the mean intensity (%) of STING staining from CTRL and TKO CD4^+^ T cells (n=7 per group). **(F)** Representative fluorescence images of cGAS (green) and dsDNA (red) in activated CD4^+^ T cells. Nuclei were counterstained with DAPI (blue). Scale bar 2 μm. **(G)** Semiquantitave analysis of the Fluorescence area of cGAS and dsDNA staining in the cytoplasm (n=4-5 per group). **(H)** Representative immunblots for Lamp1, LC3I/II and β-actin from CTRL and TKO CD4^+^ T cells in response to either DMSO or 100 nM Rapamycin for 48 hours. **(I)** Protein densitometry ratio of Lamp1/β-actin and LC3-II/β-actin by densitometry analysis (n=5 per group). **(J)** qRT-PCR showing relative mRNA expression levels of D-Loop and RNR2 from CTRL and TKO CD4^+^ T cells following DMSO or Rapamycin 100nm incubation for 48 hours. **(K)** IL-4, IL-5 and IL-13 cytokine release in activated CD4^+^ T cells following DMSO or Rapamycin 100nM incubation (n=8-12 per group). Values represent mean ± SEM. *p<0.05, **p<0.01, ***p<0.001 vs control mice by two-way ANOVA followed by the Tukey’s post hoc test or unpaired two-tailed student-t-test.

### STING activation orchestrates NF-kB activation to drive T_H_2 activation

To functionally validate that cGAS-STING activation, contributed towards NF-κB signaling and T_H_2 activation, STING was inhibited using H151 (36) or depleted via STING siRNA. H151 supplementation of primary T_H_0 CD4^+^ T cells, reduced STING levels in TKO cells (Figure 4A- B). Furthermore, the activation of IκBα, NF-κB and STAT6 were all more robustly attenuated in the TKO cells (Figure 4A-B). GATA3 levels were blunted in parallel (Figure 4A-B). Consistently, the secretion of IL-4, IL-5 and IL-13 were blunted in the TKO cells in the presence of H151 (Figure 4C). Interestingly, H151 blunted IFN-γ to a similar extent in control and TKO cells (Supplemental Figure 3A). STING knockdown (KD) was highly effective in both lineages (Supplemental Figure 3B). As STING levels were more pronounced in TKO cells, the effect of STING KD was also more robust in these CD4^+^ T cells. Here we see robust reductions STING levels, and in IκBα and NF-κB activation (Figure 4D-E). In parallel, STING KD reduced GATA3 expression to a greater extent in the TKO CD4^+^ T cells (Figure 4D-E). Similarly, STING KD signficiantly blunted IL-4, IL-5 and IL-13 secretion in the TKO cells, with a modest reduction in IL-13 secretion in control cells (Figure 4F). Similarly, and consistent with H151 supplementation, STING KD blunted IFN-γ to similar extents in control and TKO cells (Supplemental Figure 3C). The proposed mitochondrial fidelity pathway following BLOC1S1 depletion in T_H_2 cells is schematized in Figure 4G.

**Figure 4.**
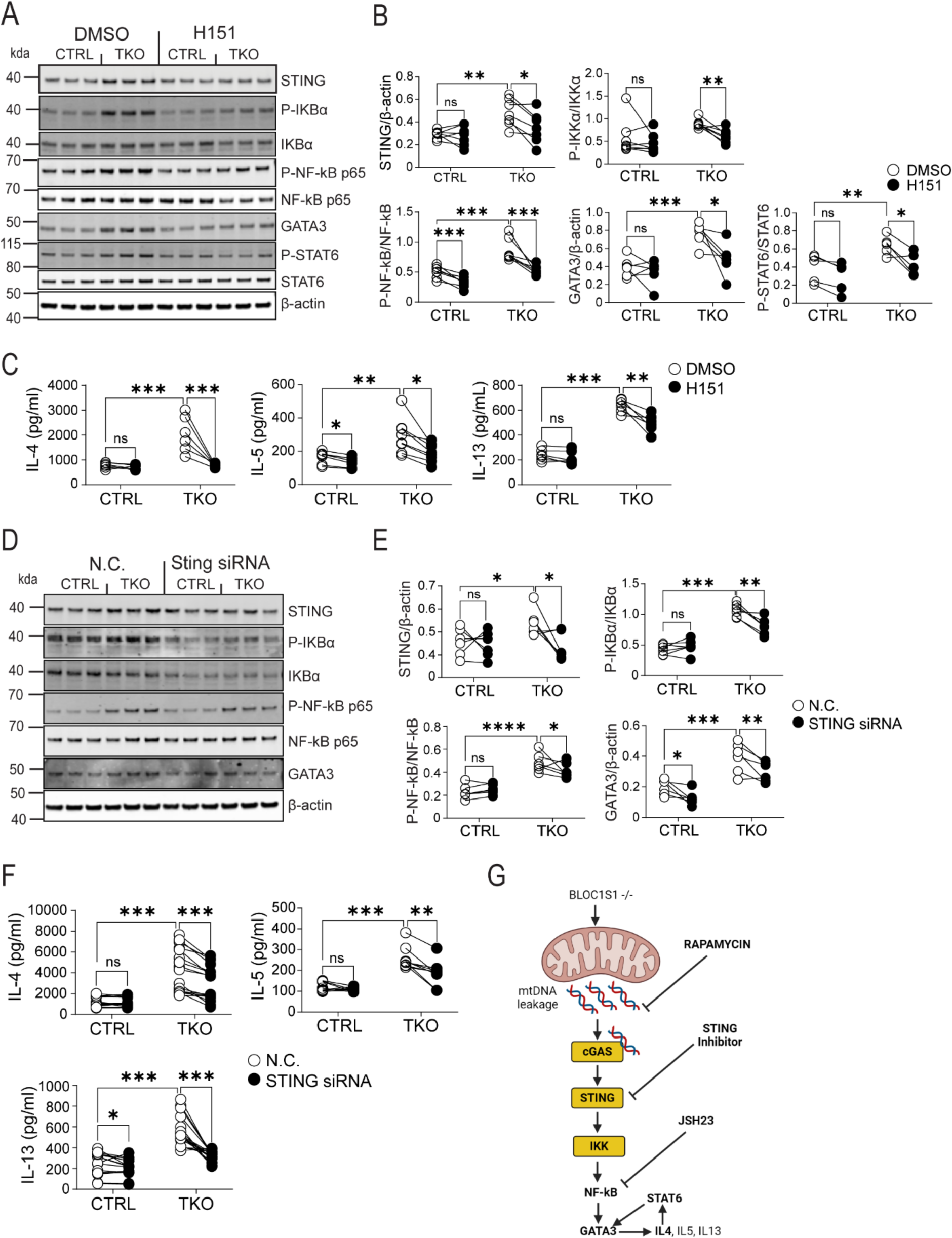
STING knockdown resulted in reduced T_H_2 cytokines in BLOC1S1-/- CD4^+^ T cells. **(A)** Representative immunoblots for STING, P-IκBα, IκBα, P-NF-κB p65, NF-κB p65, GATA3, P- STAT6, STAT6 and β-actin from CTRL and TKO CD4^+^ T cells following DMSO or H151 (500 nM) incubation for 48 hours. **(B)** Protein quantitation and ratio of STING/β-actin, P-IKBα/IKBα, P-NF-kB P65/NF-kB P65, GATA3/β-actin and P-STAT6/STAT6 by densitometry analysis (n=6-7 per group). **(C)** IL-4, IL-5 and IL-13 cytokine release in activated CD4^+^ T cells treated with either DMSO or H151 (500nM) (n=6-7 per group). **(D)** Representative immunoblots for STING, P-IκBα, IκBα, P-NF-κB p65, NF-kB p65, GATA3 and β-actin from CTRL and TKO CD4^+^ T cells incubated with negative control (N.C.) or with STING siRNA. **(E)** Protein quantitation and ratio of STING/β-actin, P-IκBα/IκBα, P-NF-κB P65/NF-κB P65 and GATA3/β-actin by densitometry analysis (n=6 per group). **(F)** IL-4, IL-5 and IL-13 cytokine release in activated CD4^+^ T cells incubated with either N.C. or STING siRNA (n=9-18 per group). **(G)** Schematic representation of proposed mechanistic pathway. Values represent mean ± SEM. *p<0.05, **p<0.01, ***p<0.001 vs control mice by two-way ANOVA followed by the Tukey’s post hoc test or unpaired two-tailed student-t-test.

### TKO mice are more susceptible to T_H_2-linked allergic disease models

As atopic disorders are driven by an elevated T_H_2 response we exploited different atopy models to functionally validate the susceptibility of T_H_2 polarization in the absence of BLOC1S1. The first model explored was atopic dermatitis (AD) induced by the topical exposure to the ear pinnae of the vitamin D3 analogue Calcipotriol (MC903) (29). The protocol is schematized in Figure 5A. This response to MC903 appeared similar in both sexes (Supplemental Figure 4A). Although, topical application of MC903 vs. ethanol control triggered an AD response with increased skin thickness in all mice, TKO mice exhibited an earlier lesion onset with more severe disease (Figure 5B). Histological examination of the red scaly lesioned skin showed greater epidermal hyperplasia with dermal lymphocyte infiltration in the TKO mice (Figure 5C and Supplemental Figure 4B). In parallel, the marker of cell proliferation, Ki67 was markedly induced in the TKO mice (Figure 5D). Consistent with atopy, TKO mice showed higher plasma levels of IgE under basal conditions, with a further exaggeration in response to the topical application of MC903 (Figure 5E). Interestingly, although BLOC1S1 deficiency was restricted to CD4^+^ T cells, qRT-PCR of whole skin samples showed that transcripts encoding bloc1s1 were blunted in this MC903 AD model in both the control and TKO mice (Figure 5F). Finally, in this model, the quantification of cytokine secretion from CD4^+^ T cells extracted from the auricular Lymph nodes showed enhanced IL-4, IL-5, and IL-13 levels in response to topical MC903 treatment in the TKO mice (Figure 5G-H).

**Figure 5.**
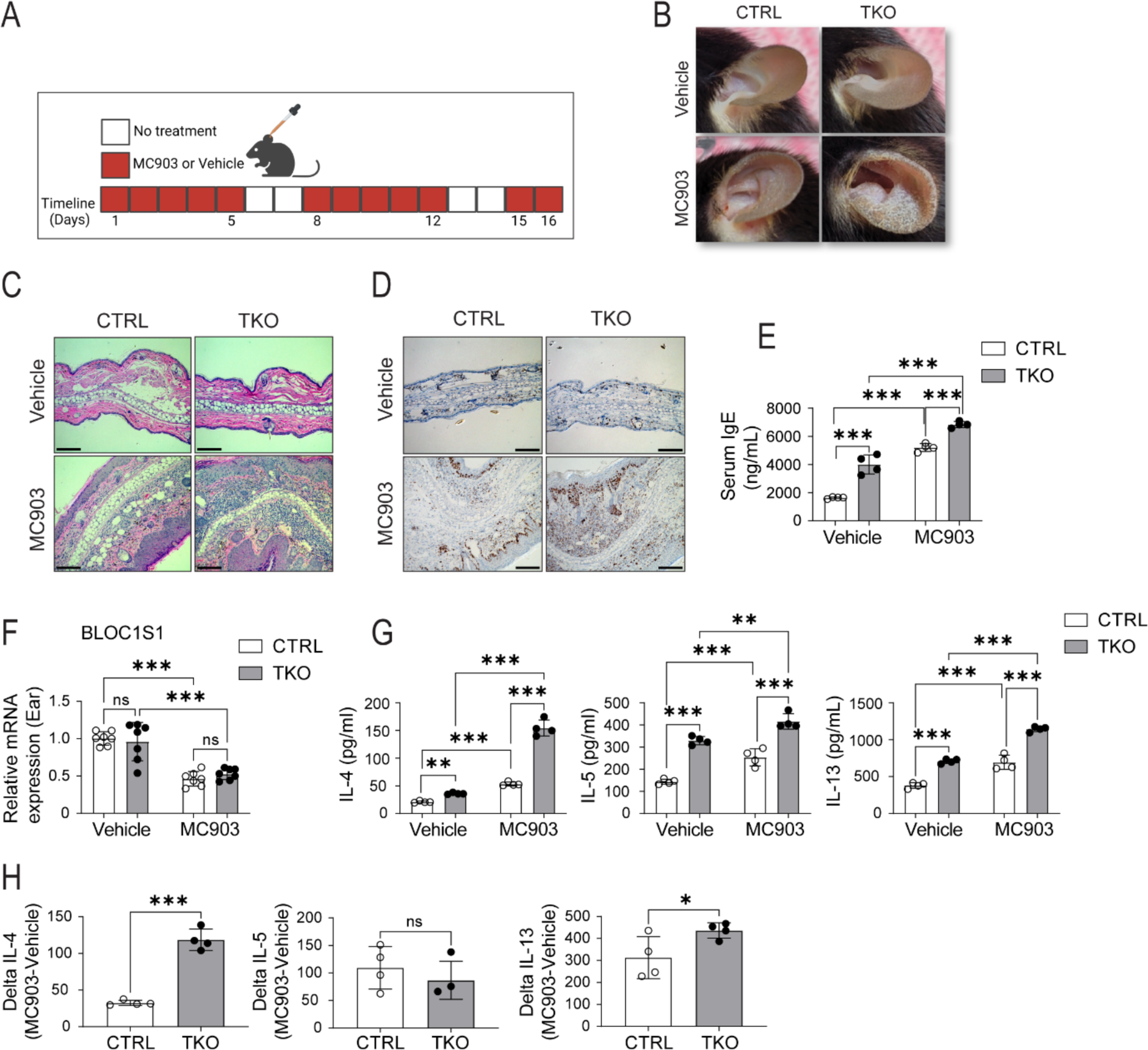
Increased calcipotriol (MC903) induced atopic dermatitis in TKO mice. **(A)** Daily study protocol for MC903 induced dermatitis (in red). **(B)** Gross appearance of Ethanol or MC903 application to CTRL and TKO mouse ears at day 12. **(C)** Representative H&E staining of ear sections at day 12. Scale bar = 100um. **(D)** Representative Ki67 staining of ear sections at day 12. Scale bar = 100um. **(E)** Plasma IgE levels from the mice following the topical application of ethanol or MC903 (n=4 per group). **(F)** qRT-PCR showing relative mRNA expression levels of BLOC1S1 from mourse ears in response to ethanol or MC903 (n=7 per group). **(G)** IL-4, IL-5 and IL-13 cytokine secreted levels at day 12 from CTRL and TKO CD4^+^ T cells isolated from auricular lymph nodes following ethanol or MC903 topical application (n=4 per group). **(H)** Delta IL-4, Delta IL-5 and Delta IL- 13 cytokine values of CTRL and TKO CD4^+^ T cells in response to MC903 (n=4 per group). Values represent mean ± SEM. *p<0.05, **p<0.01, ***p<0.001 vs control mice by two-way ANOVA followed by the Tukey’s post hoc test or unpaired two-tailed student-t-test.

We then exposed control and TKO mice to OVA-induced allergic airway inflammation as a second atopic model (30, 37). Mice were OVA challenged both systematically and via aerosol to evoke airway inflammation as depicted in Figure 6A. Histological analysis of lungs of OVA sensitized mice showed more marked peribronchial inflammation with infiltrating eosinophil infiltration in the TKO mice (Figure 6B). Consistent with a greater allergen specific type 2 response, OVA TKO mice similarly showed plasma IgE levels (Figure 6C). The TKO mice also showed greater production of IL-4 and IL-13 in CD4^+^ T cells extracted from the lungs of OVA exposed mice (Figure 6D). Furthermore, the response to OVA was relatively more robust in the TKO mice versus control mice (Figure 6E). Interestingly, and consistent with the findings in primary CD4^+^ T cell in-vitro studies, transcripts encoding STING were induced in ear from the AD model and in whole lung tissue from the allergic asthma model (Supplemental Figure 4C).

**Figure 6.**
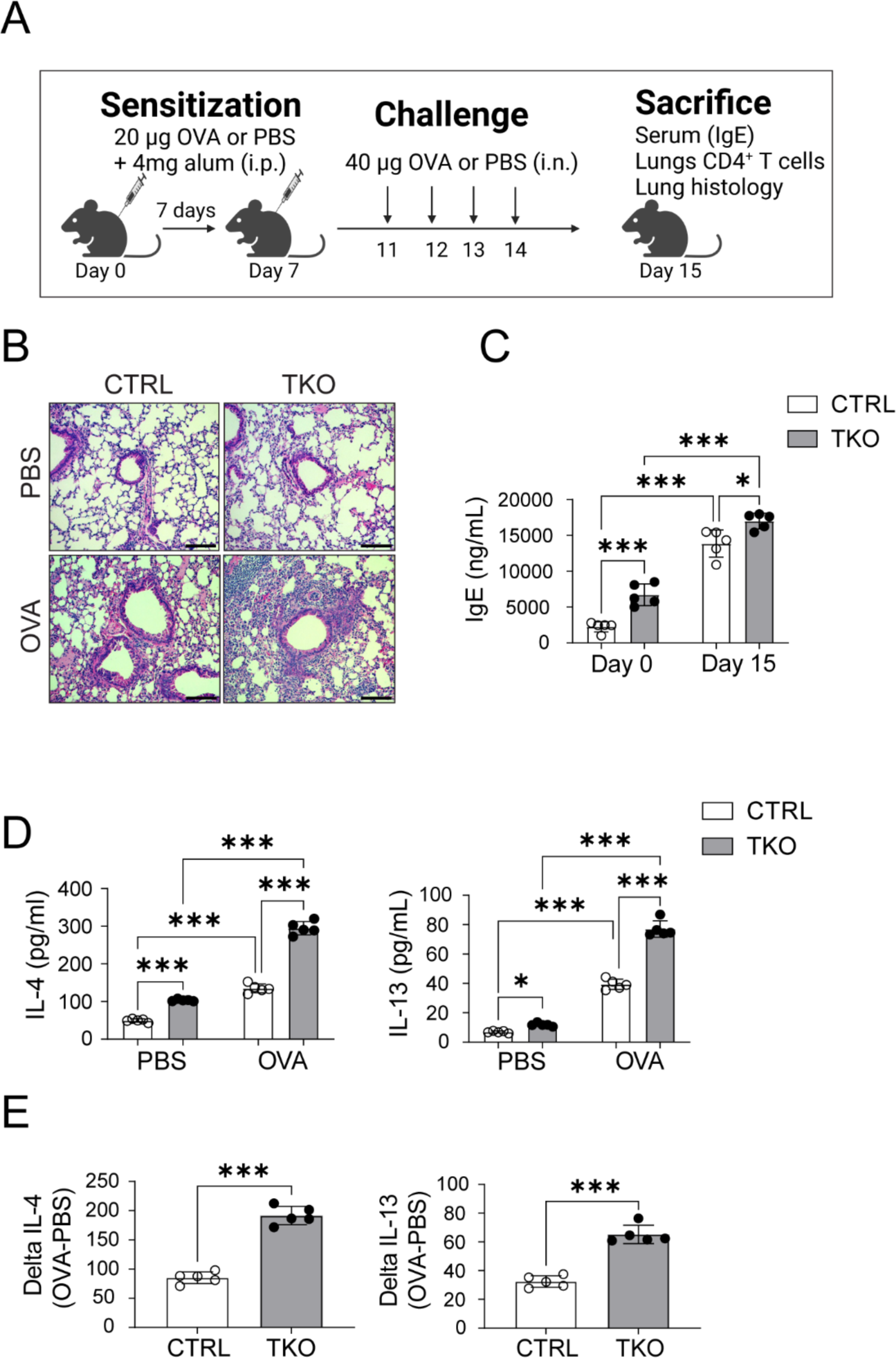
Increased ovalbumin induced airway inflammation in TKO mice. **(A)** Daily protocol for ovalbumin induced airway inflammation in mice. **(B)** Representative H&E staining of lungs of PBS and Ovalbumin (OVA) administered CTRL and TKO mice. Scale bar = 100um. **(C)** Plasma IgE from the CTRL and TKO mice following PBS or OVA administration (n=5 per group). **(D)** IL-4 and IL-13 cytokine release from CD4^+^ T cells isolated from the lungs of CTRL and TKO mice in response to PBS or OVA administration (n=5 per group). **(E)** Delta IL-4 and Delta IL-5 levels (n=5 per group). Values represent mean ± SEM. *p<0.05, **p<0.01, ***p<0.001 vs control mice by two-way ANOVA followed by the Tukey’s post hoc test or unpaired two-tailed student-t-test.

## Discussion

In this study we find that the absence of BLOC1S1 in CD4^+^ T cells, results in the preferential polarization into the T_H_2 cell lineage in response to TCR engagement. This programing appears to be driven by perturbed mitochondrial fidelity possibly coupled to incomplete autophagic clearance of mtDNA. The persistently increased cytosolic mtDNA, activates the cGAS-STING immune surveillance program. Downstream of this, NF-κB activation is linked with the induction of GATA3 and phosphorylated STAT6 to amplify T_H_2 lineage responsiveness. Consistent with this immune phenotype, the BLOC1S1 TKO mice show greater susceptibility to allergic conditions including atopic dermatitis and allergic asthma.

The cell autonomous program promoting T_H_2 polarization is well established with respect to the role of GATA3 and STAT6 signaling. However, the roles of mtDNA initiated signaling and cGAS-STING activation has not been well established in adaptive immune cells. In contrast, in innate immunity, mtDNA functions as a canonical damage associated molecular pattern (DAMP) to initiate cGAS-STING signaling myeloid cells (34), and to drive neutrophil NETosis (38). Recent data does show that aging is linked with the disruption of lysosome proteasomal function with a concomitant increase in mtDNA and inflammaging in human CD4^+^ T cells (39). Although, the role of cGAS-STING or NF-κB was not assessed in that study (39). Conversely, in the murine tumor microenvironment CD4^+^ T-intrinsic STING activation drives T_H_1 and T_H_9 activation (40). Interestingly here, the T_H_1, but not the T_H_9 phenotype was dependent on type 1 interferon signaling, although both lineages were dependent on MTOR and NF-κB activity (40). At the same time, the genetic disruption of NF-κB signaling attenuated allergic airway induced T_H_2 polarization (30). Together these data highlight that the fate of distinct CD4^+^ T cell subsets have different signaling pathways, that may be moderated in part by the in-situ environment and immune cell cross talk. Our study adds to these finding by showing the depleting BLOC1S1 initiates a mtDNA, cGAS-STING and NF-κB integrated pathway that preferentially augments T_H_2 cell responsiveness and susceptibility to atopy.

In innate immunity, TBK1 is a canonical kinase in the STING mediated induction of NF-kB too enable signal transduction (41). This STING mediated activation of TBK1 results in the subsequent phosphorylation of NF-κB essential modulator (NEMO), a regulatory subunit of the IKK (inhibitor of NF-κB kinase) complex (42). Phosphorylated NEMO then activates the IKK complex, leading to the degradation of inhibitory IkB proteins and the release and nuclear translocation of active NF-κB (42). In parallel, TBK1 activation promotes degradation of STING via the induction of autophagy as a negative regulatory feedback loop (43). Interestingly, the absence of BLOC1S1 is known to constipate the autophagolysosomal degradation pathway (24), and whether this pathway is operational in augmented cytosolic mtDNA and or accumulation of STING in the TKO cells warrants further evaluation. Contrary to the effects on innate immunity, the conditional knockout of TBK1 in CD4^+^ T cells resulted in the induction of IFN-γ and of CD4^+^ memory cells, and the pharmacologic inhibition of TBK1 blunted experimental autoimmune encephalitis (44). Conversely our study shows the TKO mice exhibit an increased T_H_2 profile with the exacerbation of allergic disease in parallel with increased TBK1 phosphorylation. Although these data remain to be reconciled, this may point to the specific roles of BLOC1S1 in mitochondrial quality control and/or in autophagosome homeostatic functions to alleviate cytosolic DAMPS and thereby reduce T_H_2 polarization.

The concept that intracellular quality control programs moderate immune responsiveness is well established in programs controlling autophagy and mitochondria homeostasis. Emerging studies are implicating immunoregulatory effects of programs controlling lipid handling and endo-lysosomal function. At a reductionist levels, these distinct programs are being well characterized, and at a systemic biology perspective, these programs play integrated roles. Interestingly, BLOC1S1 as a nutrient-sensing homeostatic mediator contribute towards the control of all these programs (15, 25). In this manuscript, we show that the absence of BLOC1S1 has a preferential effect on enhancing T_H_2 immune cell responsiveness. The comprehensive mechanisms of action remain to be determined. However, our initial findings support both that the extrusion of mtDNA from mitochondria and the accumulation of STING on endo-lysosomes which together may amplify both canonical and non-canonical T_H_2-linked immune responsiveness.

Genetic defects in BLOC1S1 have been associated with juvenile leukodystrophy (45) and has recently been linked with a reduction in lysosome content and increased lipid stores (26). Although juvenile leukodystrophy is associated with neuroinflammation, a direct link to T_H_2 biology does not appear to have been explored (46). Furthermore, lipid biology is emerging as a mediator pathway in CD4^+^ T cell polarization (47). At the same time, it is interesting that transcriptomic analysis in 16 atopic dermatitis subjects, shows that BLOC1S1 expression is significantly blunted in association with this allergic ichthyosis (48). Given that human iPSC’s are available that incorporate BLOC1S1 mutations (49), these cells have the potential for the direct exploration of the role of BLOC1S1 genetic mutations in CD4^+^ T cell biology.

In conclusion, we show that in the absence of BLOCS1, which exhibits perturbed mitochondrial and endolysosomal functioning, that CD4^+^ T cells are preferentially polarization towards the T_H_2 lineage. This is shown to be mediated in part, but cytosolic mtDNA accumulation, STING activation with downstream NFkB signaling, which drives the canonical GATA3 and STAT6 signaling to augment T_H_2 responsiveness. This T_H_2 signature is validated in vivo with increased atopic dermatitis and allergic asthma in the BLOC1S1 knockout mice. This model should allow us to further dissect out the integrated roles of mitochondrial and endolysosomal homeostasis programs in driving CD4^+^ T cell fates.

## Acknowledgements

This research was supported by the NHLBI Division of Intramural Research (MNS – ZIA- HL005199). We thank and acknowledge the assistance of the NHLBI Laboratory Animal Core Facility. Finally, we thank Nina Boehm (née Klimova) Ph.D., who initiated this project in the laboratory.

