## Supplementary Material for "BLOC1S1 control of vacuolar organelle fidelity modulates T_H_2 cell immunity and allergy susceptibility"

#### TITLE

**Suppl. Figure 1.** BLOC1S1 depleted $CD4^+$ T cells preferentially augments $T_H2$ immune response.

**Suppl. Figure 2.** Increased $Lamp1^+$ Cells in $BLOC1S1^{-/-}$ $CD4^+$ T-cells.

**Suppl. Figure 3.** STING inhibition and its siRNA KD reduces IFN-$\gamma$ levels in CTRL and $BLOC1S1^{-/-}$ $CD4^+$ T-cells.

**Suppl. Figure 4.** TKO mice are more susceptible to drug induced dermatitis than CTRL mice.

****Suppl. Table 1.** List of primers used for qRT-PCR analysis.

****Suppl. Table 2.** List of antibodies used for immunofluorescence and immunoblotting.

****Suppl. Table 3.** List of FACS antibodies used and panel design for FACS analysis.

****Suppl. Table 4.** List of reagents used.

#### Supplementary Figure 1

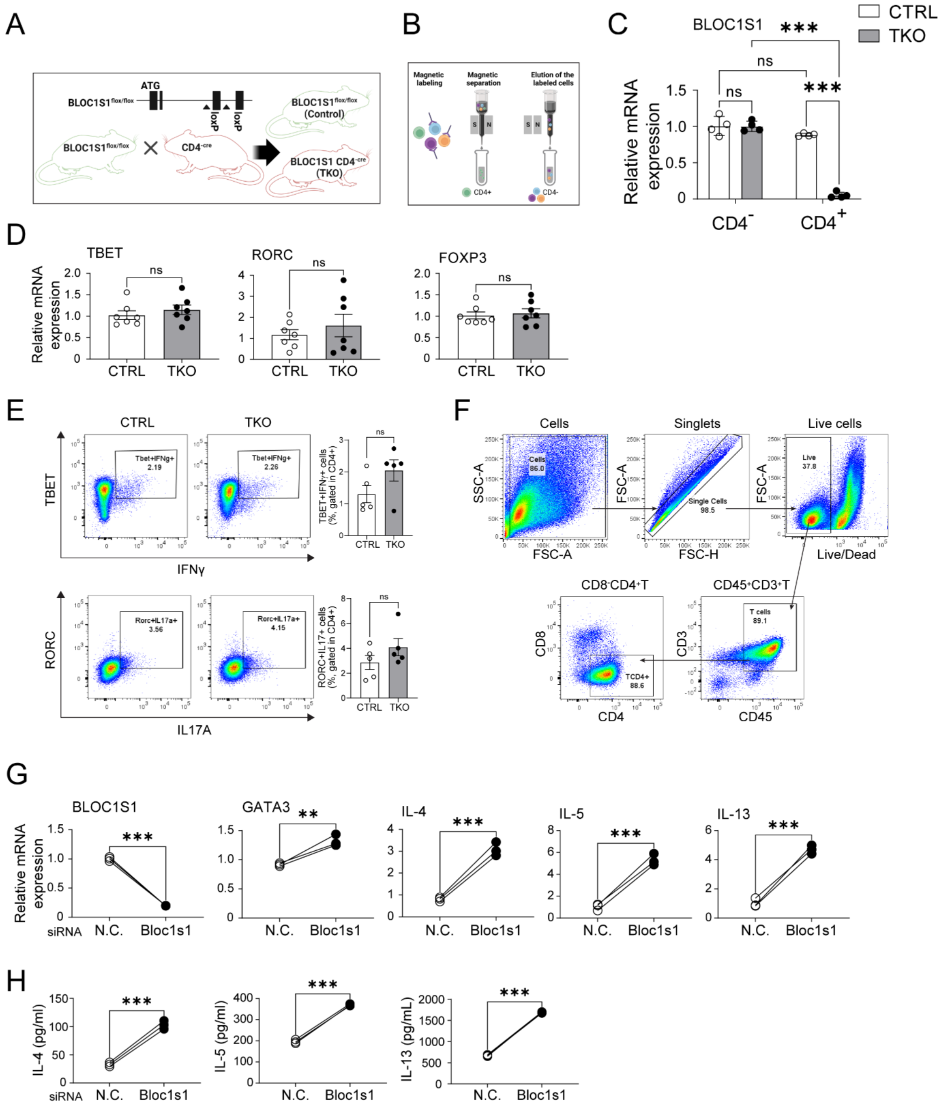

**Figure S1. BLOC1S1 depleted CD4$^+$ T cells preferentially augments $T_H2$ immune response.**

**(A)** Schematic representation of approach to generate CD4$^+$ cell specific BLOC1S1 knockout (TKO) mouse. **(B)** Schematic representation of approach to separate CD4$^+$ T cells from the residual splenic pool (CD4$^-$ cells). **(C)** qRT-PCR showing relative mRNA expression levels of BLOC1S1 in CD4$^+$ and CD4$^-$ cells (n=4 group). **(D)** qRT-PCR showing relative mRNA expression levels of TBET, RORC and FOXP3 in CD4$^+$ T cells (n=7 per group). **(E)** Representative flow-cytometric analysis of intracellular cytokines TBET$^+$IFN-γ$^+$ and RORC$^+$IL17$^+$ in CD4$^+$ T cells (n=5 per group). **(F)** Representative flow-cytometry gating profile of CD4$^+$ T cells for flow cytometry. **(G)** qRT-PCR showing relative mRNA expression levels of BLOC1S1, GATA3, IL-4, IL-5 and IL-13 in CD4$^+$ T cells isolated from blood of healthy individuals treated with either N.C. or siRNA (n=3 individuals). **(H)** IL-4, IL-5 and IL-13 cytokine release in activated CD4$^+$ T cells isolated from blood of healthy individuals (n=3 per group). Values represent mean ± SEM. *p<0.05, **p<0.01, ***p<0.001 vs. control mice using unpaired two-tailed student-t-test. FSC, forward scatter; SSC, side scatter.

#### Supplementary Figure 2

Commented [HKK([1]: It's too big? Please adjust it with same size of other figures

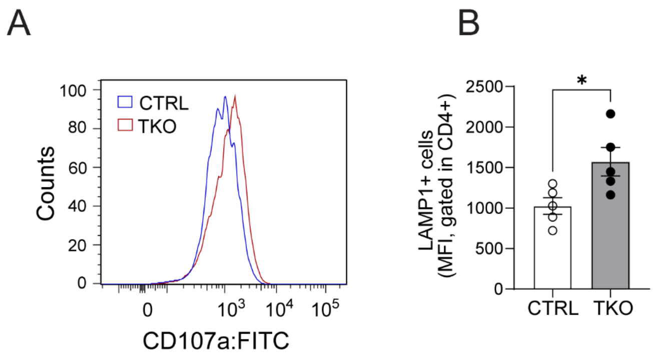


**Figure S2. Increased Lamp1$^+$ Cells in BLOC1S1$^{-/-}$ CD4$^+$ T cells.**

**(A)** Intracellular staining of CTRL and TKO mouse CD4$^+$ T cells with CD107a (Lamp-1). **(B)** Histogram of Lamp-1$^+$ in CD4$^+$ T cells (n=5 per group). Values represent mean +/- standard error of mean (SEM). *p<0.05, **p<0.01, ***p<0.001 vs. CTRL by using unpaired two-tailed student-t-test.

### Supplementary Figure 3

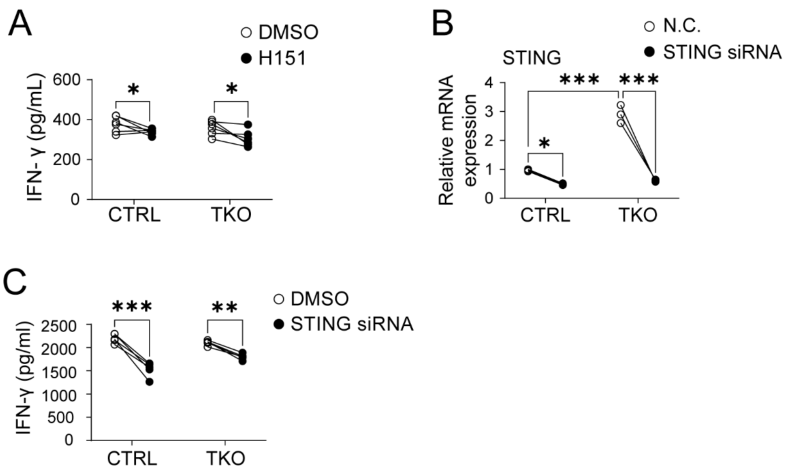


**Figure S3. STING inhibition and its siRNA KD reduces IFN-γ levels in CTRL and BLOC1S1$^{-/-}$ CD4$^{+}$ T cells.**

**(A)** IFN-γ cytokine release in CTRL and TKO CD4$^{+}$ T cells treated with either DMSO or H151 (500 nM) (n=6 per group). **(B)** qRT-PCR showing relative mRNA expression levels of STING in CD$^{+}$ T cells of CTRL and TKO treated with either N.C. or STING siRNA (n=3 per group). **(C)** IFN-γ cytokine release in CTRL and TKO CD4$^{+}$ T cells treated with either N.C. or STING siRNA (n=6 per group). Values represent mean ± SEM. *p<0.05, **p<0.01, ***p<0.001 vs control mice by two-way ANOVA followed by the Tukey's post hoc test or unpaired two-tailed student-t-test.

#### Supplementary Figure 4

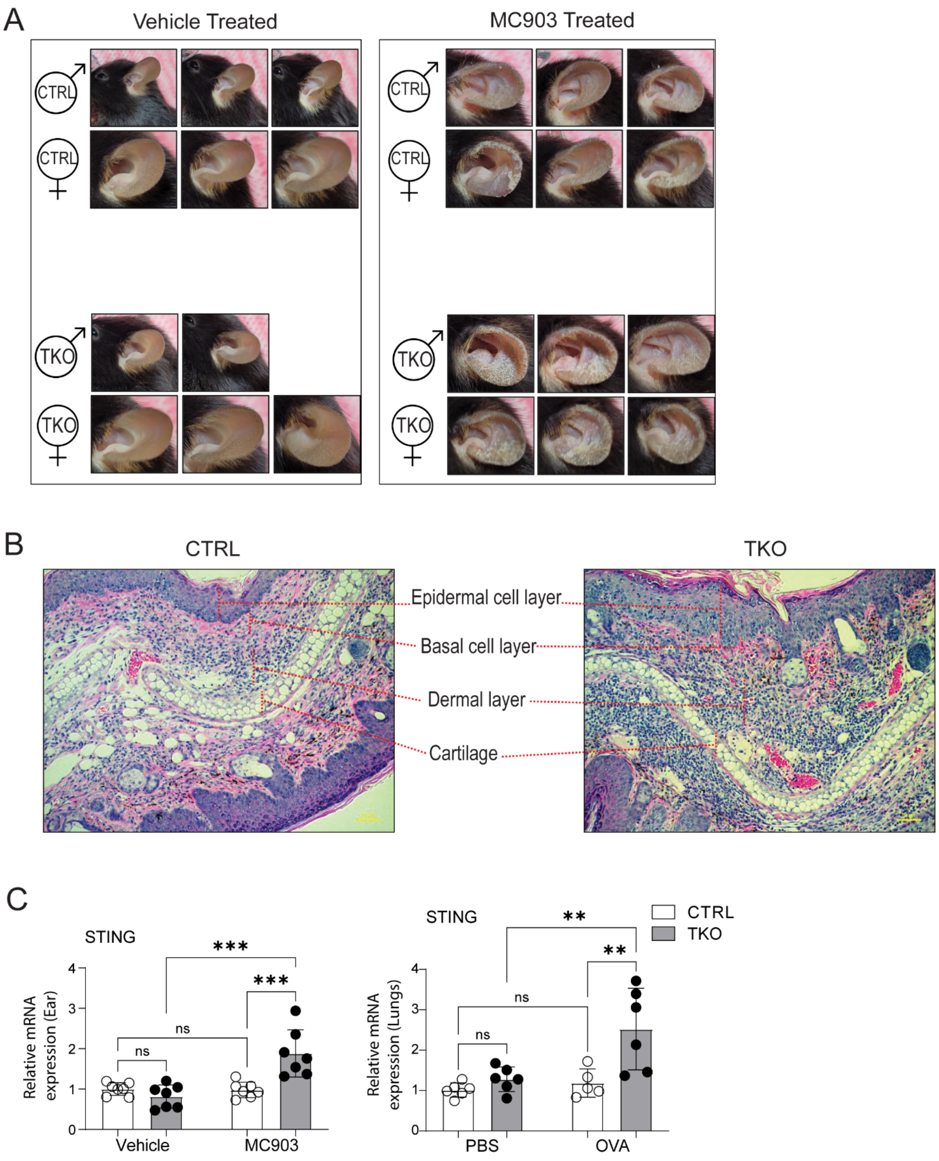

**Figure S4. TKO mice are more susceptible to drug induced dermatitis than CTRL mice.**

**(A)** Gross appearance of ears of mice treated with either ethanol or MC903 at day 12 in both sexes. **(B)** Relative H&E staining of MC903 treated CTRL and TKO mice ear sections at day 12. **(C)** qRT-PCR showing relative mRNA expression levels of STING in MC903 treated ear and OVA treated lungs (n=5-7 per group). Values represent mean $\pm$ SEM. *p<0.05, **p<0.01, ***p<0.001 vs. control mice by two-way ANOVA followed by the Tukey's post hoc test or unpaired two-tailed student-t-test.

**Supplementary Table 1.**

Sequence of primers used for qRT-PCR studies.

**Commented [HKK([2]:** Table line space and style looks different
As well as please use same style of table

| Name | Source | Oligonucleotides |
|---|---|---|
| IL-4 (Mouse) | Integrated DNA Technologies | F: CAAACGTCCTCACAGCAACG<br>R: TGCAGCTCCATGAGAACACTAG |
| IL-5 (Mouse) | Integrated DNA Technologies | F: AGCAATGAGACGATGAGGCTTC<br>R: CCCACGGACAGTTTGATTCTTCAG |
| IL-13 (Mouse) | Integrated DNA Technologies | F: AAGATCTGTGTCTCTCCCTCTGAC<br>R: ATACCATGCTGCCGTTGCAC |
| RNR2 (Mouse) | Integrated DNA Technologies | F: CTAGAAACCCCGAAACCAAA<br>R: CCAGCTATCACCAAGCTCGT |
| D-Loop (Mouse) | Integrated DNA Technologies | F: AATCTACCATCCTCCGTGAAACC<br>R: TCAGTTTAGCTACCCCCAAGTTTAA |
| TERT (Mouse) | Integrated DNA Technologies | F: CTAGCTCATGTGTCAAGACCCTCTT<br>R: GCCAGCACGTTTCTCTCGTT |
| BLOC1S1 (Mouse) | Integrated DNA Technologies | F: TCCCGCCTGCTCAAAGAAC<br>R: GAGGTGATCCACCAACGCTT |
| FoxP3 (Mouse) | Integrated DNA Technologies | F: CACCCAGGAAAGACAGCAACC<br>R: GCAAGAGCTCTTGTCCATTGA |
| T-bet (Mouse) | Integrated DNA Technologies | F: TCAACCAGCACCAGAGAGAG<br>R: AAACATCCTGTAATGGCTTGTG |
| Rorc (Mouse) QuantiTech Primer | Qiagen | Cat. QT00197722 |
| 18S (Mouse) QuantiTech Primer | Qiagen | Cat. QT02448075 |

| | | |
|---|---|---|
| 16S (Mouse) | Integrated DNA Technologies | F: CTAGAAACCCCGAAACCAAA<br>R: TCAGTTTAGCTACCCCCAAGTTTAA |
| IL-4 (Human) QuantiTech Primer | Qiagen | Cat. QT00012565 |
| IL-5 (Human) QuantiTech Primer | Qiagen | Cat. QT00001435 |
| IL-13 (human) | Qiagen | Cat. QT00000511 |
| GATA3 (Human) | Integrated DNA Technologies | F: GAACCGGCCCCTCATTAAG<br>R: ATTTTTCGGTTTCTGGTCTGGAT |
| BLOC1S1 (Human) QuantiTech Primer | Qiagen | Cat. QT0016002 |
| 18S (Human) QuantiTech Primer | Qiagen | Cat. QT00199367 |

**Supplementary Table 2.**

| Antibodies used for immunoblotting and Immunofluorescence | | | |
|---|---|---|---|
| **Antibody** | **Catalog #** | **Working dilution** | **Source** |
| Ki67 (D3B5) | 9129S | 1:400 (IF) | Cell Signaling |
| P-IKB-alpha (Ser32) (14D4) | 2859S | 1:1000 (IB) | Cell Signaling |
| IKB-alpha | 9242S | 1:1000 (IB) | Cell Signaling |
| P-NFkB P65 (Ser536) (93H1) | 3033S | 1:1000 )IB) | Cell Signaling |
| NFkB P65 (D14E12) | 8242S | 1:1000 (IB) | Cell Signaling |
| P-STAT6 (Tyr641) (D8S9Y) | 56554S | 1:1000 (IB) | Cell Signaling |
| STAT6 (D3H4) | 5397S | 1:1000 (IB) | Cell Signaling |
| GATA3 (D13C9) | 5852S | 1:1000 (IB) | Cell Signaling |
| β-actin (8H10D10) | 3700S | 1:1000 (IB) | Cell Signaling |
| Total Oxphos | Ab110413 | 1:1000 (IB) | Abcam |
| cGAS (D3080) | 31659S | 1:1000 (IB) | Cell Signaling |
| P-STING (Ser365) (D8F4W) | 72971S | 1:1000 (IB) | Cell Signaling |
| STING (D2P2F) | 13647S | 1:1000 (IB), 1:200 (IF) | Cell Signaling |
| P-TBK1 (Ser172) (D52C2) | 5483S | 1:1000 (IB) | Cell Signaling |
| TBK1 (E813G) | 38066S | 1:1000 (IB) | Cell Signaling |
| Tom20 (F-10) | SC-17764 | 1:1000 (IB), 1:200 (IF) | Santa Cruz |
| VDAC (D73D12) | 4661S | 1:1000 (IB) | Cell Signaling |
| Lamp1 (ab208943) | Ab208943 | 1:1000 (IB), 1:200 (IF) | Abcam |
| LC3 AB (D3U4C) | 12741S | 1:1000 (IB), 1:200 (IF) | Cell Signaling |
| dsDNA (HYB331-01) | SC-58749 | 1:200 (IF) | Santa Cruz |
| Lamp1 (H4A3) | SC-20011 | 1:1000 (IB), 1:200 (IF) | Santa Cruz |

**Supplementary Table 3.**

Antibody panel design for flow cytometry.

| Target | Version | Catalog# | Vendor |
|---|---|---|---|
| CD4 | BUV 395 | 563790 | BD |
| CD3 | BUV496 | 741117 | BD |
| IFN-g | BUV737 | 612769 | BD |
| CD45 | BUV805 | 568336 | BD |
| LIVE_DEAD | BV421 | 423114 | BIOLEGEND |
| TNF-a | BV510 | 506339 | BIOLEGEND |
| CD8 | BV605 | 563152 | BD |
| RORGT | BV650 | 564722 | BD |
| IL-4 | BV711 | 504133 | BIOLEGEND |
| IL-17A | BV786 | 564171 | BD |
| CD107A | FITC | 121606 | BIOLEGEND |
| CD27 | PERCPCY5.5 | 563603 | BD |
| IL-13 | PE | 568551 | BD |
| TBET | PECF594 | 562467 | BD |
| MHC-II | PECY7 | 107630 | BIOLEGEND |
| FOXP3 | APC | 567462 | BD |
| GATA3 | APC-700 | 567633 | BD |
| CD44 | APCCY7 | 103028 | BIOLEGEND |
| | FOXP3 BUFFER | 00-5523-00 | E BIOSCIENCE |
| | leukocyte activation cocktail | 550583 | BD |

**Supplementary Table 4.**

List of reagents.

| Reagent | Catalog number | Source |
|---|---|---|
| CyQuant Cell Proliferation Assay | C7026 | Invitrogen |
| Pierce BCA Protein Assay | 23227 | Thermo Scientific |
| Human IL-4 Duoset ELISA Kit | DY204 | R&D Systems |
| Human IL-5 Duoset ELISA Kit | DY205 | R&D Systems |
| Human IL-13 Duoset ELISA Kit | DY213 | R&D Systems |
| Mouse IFN-gamma Duoset ELISA Kit | DY485 | R&D Systems |
| Mouse TNF-alpha Duoset ELISA Kit | DY410 | R&D Systems |
| Mouse IL-4 Duoset ELISA Kit | DY404 | R&D Systems |
| Mouse IL-5 Duoset ELISA Kit | DY405 | R&D Systems |
| Mouse IL-13 Duoset ELISA Kit | DY413 | R&D Systems |
| Mouse IL-10 Duoset ELISA Kit | DY417 | R&D Systems |
| Mouse IL-17 Duoset ELISA Kit | DY421 | R&D Systems |
| Lymphocyte Separation Medium | 0850494 | MP Biomedicals |

| | | |
|---|---|---|
| Human CD4+ T Cell Isolation Kit | 130-096-533 | Miltenyi Biotec |
| NucleoSpin RNA Kit | 740955 | Macherey-Nagel |
| First-strand Synthesis SuperMix | 11752250 | Invitrogen |
| FastStart Essential DNA Green Master Mix | 06924204001 | Roche Life Science |
| Accell siRNA delivery media | B-005000-100 | Horizon Discovery |
| H151 | HY-112693 | MedChemExpress |
| JSH-23 | HY-13982 | MedChemExpress |
| Rapamycin | HY-10219 | MedChemExpress |
| SMARTpool: Accell BLOC1S1 | E-012580-00-0020 | Dharmacon |
| SMARTpool: Accell Tmem173 | E-055528-00-0020 | Dharmacon |
| CD3 (Mouse) | 100339 | Biolegend |
| CD28 (Mouse) | 102116 | Biolegend |
